# Maintenance of pig brain function under extracorporeal pulsatile circulatory control (EPCC)

**DOI:** 10.1101/2023.05.28.542663

**Authors:** Muhammed Shariff, Aksharkumar Dobariya, Obada Albaghdadi, Jacob Awkal, Hadi Moussa, Gabriel Reyes, Mansur Syed, Robert Hart, Cameron Longfellow, Debra Douglass, Tarek Y. El Ahmadieh, Levi B. Good, Vikram Jakkamsetti, Gauri Kathote, Gus Angulo, Qian Ma, Ronnie Brown, Misha Dunbar, John M. Shelton, Bret M. Evers, Sourav Patnaik, Ulrike Hoffmann, Amy E. Hackmann, Bruce Mickey, Matthias Peltz, Michael E. Jessen, Juan M. Pascual

**Author notes:** **Correspondence:** Juan M. Pascual, MD, PhD, The Once Upon a Time Foundation Professor in Pediatric Neurologic Diseases, Ed and Sue Rose Distinguished Professor in Neurology, Director, Rare Brain Disorders Program, UT Southwestern Medical Center, 5323 Harry Hines Blvd. Mail code 8813, Dallas, TX 75390-8813, |. Equal contribution (Muhammed Shariff, Aksharkumar Dobariya, Obada Albaghdadi). Department of Neurosurgery, Loma Linda University Medical Center, Loma Linda, CA, 92354, USA. Heart and Vascular Center Brigham and Women’s Hospital, Boston, MA 02115, USA.

## Abstract

Selective vascular access to the brain is desirable in metabolic tracer, pharmacological and other studies aimed to characterize neural properties in isolation from somatic influences from chest, abdomen or limbs. However, current methods for artificial control of cerebral circulation can abolish pulsatility-dependent vascular signaling or neural network phenomena such as the electrocorticogram even when preserving individual neuronal activity. Thus, we set out to mechanically render cerebral hemodynamics fully regulable to replicate or modify native pig brain perfusion. To this end, blood flow to the head was surgically separated from the systemic circulation and full extracorporeal pulsatile circulatory control (EPCC) delivered via a modified aorta or brachiocephalic artery. This control relied on a computerized algorithm that maintained, for several hours, blood pressure, flow and pulsatility at near-native values individually measured before EPCC. Continuous electrocorticography and brain depth electrode recordings were used to evaluate brain activity relative to the standard offered by awake human electrocorticography. Under EPCC, this activity remained unaltered or minimally perturbed compared to the native circulation state, as did cerebral oxygenation, pressure, temperature and microscopic structure. Thus, our approach enables the study of neural activity and its circulatory manipulation in independence of most of the rest of the organism.

## 1. Introduction

Cerebral blood flow manipulation enables or facilitates numerous experimental approaches such as the investigation of brain function under ischemia and stroke or the study of the action and distribution of pharmacological agents or metabolic tracers ^1^. In some of these studies, selective vascular access to head or brain is desirable to characterize neural properties as independently as possible from somatic influences stemming from unavoidable native or experimentally induced changes in the rest of the organism. This is because metabolism or multi-organ distribution of biologically active substances can present insurmountable limitations to accurate kinetic analyses ^2^. Among these influences, blood flow and pressure alterations and extracerebral metabolism can impose significant and often nonlinear effects on brain function. To circumvent these limitations, we reasoned that the simplest approach would entail vascular disconnection followed by artificial provision of the mechanical, cellular and chemical circulatory support and control that the rest of the body exerts on the brain. A subsequent objective was to enable, via this regulation, the future investigation of a broad variety of physiological and pathological circulatory states. To this end, we built a computer-controlled mechanical perfusion and blood composition regulatory system suited for direct coupling to the arterial and venous circulation of the pig. We refer to this system as extracorporeal pulsatile circulatory control (EPCC).

Artificial perfusion of large animal brains has been practiced for decades using a variety of techniques. In these settings, monkey ^3, 4^ and canine brains ^5, 6^ have displayed retention of rudimentary neurophysiological function. Specifically, these studies have generally yielded only markedly suppressed activity. Similarly, a postmortem pig brain model subject to mechanical perfusion illustrates that even when perfusion allows for the maintenance of an extensive array of local and individual neuronal activities, electrocorticography (ECoG) is abolished ^7^. As the summation of innumerable cellular activities and connectivity highly organized into oscillatory and other periodic signals that change over well-defined time intervals, ECoG represents a robust measure of neural function integrity.

Thus, our primary objective was to preserve brain function under EPCC to achieve circulatory isolation from the majority of the rest of the body in fully controllable fashion. Brain function was measured under minimal anesthesia interference ^8^ and inferred from the integrity of high-order network activity as reported by the quantification of broad-spatial scale electrical field potential recordings from the surfaces and depths of the brain. To enable neurophysiological studies faithful to native recorded conditions, we aimed to preserve higher order neural activity as reported by electrocorticography and field potential intracerebral depth recordings. Both types of recording are vulnerable within seconds or minutes to alterations in perfusion as attested by clinical and experimental experience ^9 10^. We considered these recordings a standard of functional activity because the encephalogram is highly susceptible to, and changes in specific fashion upon, alterations in blood pressure ^11^. Thus, to assess preservation of cerebral activity under EPCC, we obtained nearly continuous, individualized pre- and post-EPCC recordings to evaluate neurophysiological impact. This showed negligible or no changes after EPCC. In addition to performing these pre and post EPCC comparisons, we also compared ECoG and depth recordings pre and post EPCC with independent pig recordings obtained under no EPCC, and evaluated pig ECoG pre and post EPCC in relation to our awake human ECoG recordings ^8^ and also found them essentially indistinguishable (except for the segment of ECoG activity related to open eyes).

Of note, the goal was to develop a method capable of preserving perfusion and neurophysiological activity at any native (i.e., pre-EPCC) individual animal value, rather than to modify perfusion to approximate a normative or uniform group value in several animals. For the question asked was whether, for any one animal, our EPCC system allowed the parameters of interest to remain constant or near-constant relative to the pre-EPCC state.

## 2. Materials and methods

### 2.1 Conceptualization and general design of EPCC

Taking some of the limitations of previous approaches to the perfusion of the brain and other organs into account, EPCC was guided by four physiological considerations.

First, we opted for pulsatile instead of linear flow. Even though today’s clinical extracorporeal perfusion is usually synonymous with non-pulsatile circulation, pulsatility has been recommended to maintain human extracerebral organ function in medical practice ^12^. Yet, clinical practice or studies provide a limited rationale for pulsatility partly because their observations are, by necessity, heterogeneous or incompletely descriptive of the rich physiology of brain perfusion. Instead, more systematic reasons stem from animal and cellular observations. On the one hand, pulsatility impacts, at least in experimental animal pathophysiological states, regional and global cerebral blood flow (CBF), cerebral metabolic rate of oxygen (CMRO_2_), cerebral delivery of oxygen (CDO_2_) and cerebral vascular resistance (CVR) ^13^, and this may exert neural function repercussions since the recovery from global brain ischemia can be influenced by pulsatility ^14^. On the other, despite uncertainties about specific cell signaling mechanisms, cerebral endothelial cells ^15^, pericytes ^16^ and astrocytes ^17^ are baroreceptive and exhibit heartbeat frequency-range responsiveness. This phenomenon is associated with the regulation of molecules relevant to neurovascular coupling and other salient facets of neural activity. For example, nitric oxide synthase activity is markedly reduced under nonpulsatile flow, limiting vasodilation capacity in addition to increasing the risk for thrombosis in relation to abnormal platelet aggregation ^18^. Thus, we reasoned that pulsatility would minimize or avoid disruption of potentially significant biological mechanisms.

Second, given the normal variability of circulatory function across pigs, which is also noted in man and other animal research models, and the potential for further increase of this variability under experimental interventions ^19^, we performed individualized measurements prior to EPCC in each pig. This allowed us to customize EPCC to maintain circulation to the brain as closely as feasible to the native state of each animal following a negligible delay upon vascular diversion, thus accounting for pre-EPCC individual variability or abnormalities. This also implied that, to fulfill the purposes of this study, we did not manipulate native blood pressure to meet any particular pre-determined or normative values. Instead, we accepted any native values and tested their replicability by EPCC.

Third, to render EPCC useful for a variety of potential experimental situations, we aimed for broad modulatory capacity of blood pressure, pulsatility, flow and as many cellular and chemical composition parameters as desirable in future studies. Thus, we designed EPCC with variable circulatory inputs (such as systolic and diastolic pressure, flow or beat rate) that could be adjusted, and measured and manipulated blood composition as needed to maintain stability when so desired. The testing of these modifiable capabilities beyond the maintenance of a stable, near-native circulation was not a goal of this study and will require further work.

Fourth, although blood substitutes are available, we retained each pig’s autologous blood and supplemented it with heterologous whole blood. This aimed to minimize chemical composition changes while maintaining red blood cell and brain metabolic relations ^20^.

### 2.2 Animals

This study was approved by the Institutional Animal Care and Use Committee of UT Southwestern Medical Center. All other relevant institutional regulations were followed, in addition to ARRIVE (Animal Research: Reporting of In Vivo Experiments) guidelines. Domestic farm pigs (*Sus scrofa domesticus*) were studied. In addition to the pigs from which the data presented below were obtained, other groups of pigs were used for preparative work and supportive aspects. These groups are listed here to facilitate the full reproducibility of the experimental context. First, preliminary studies were conducted in cumulatively modified fashion, in 4 anesthetized juvenile pigs of 4 - 5 months of age weighing 20 - 25 Kg. This weight was chosen because of progressive limitation to surgical intracranial accessibility with age and because of near completion of brain growth at 5 months of age ^21^. These pigs were used to develop isolated aspects of surgery, angiography or EPCC and therefore are not individually described besides the fact that the knowledge gained during this preliminary work is reflected in the methods presented here. Second, 6 pigs weighing 30 – 60 Kg used in unrelated terminal studies were exsanguinated under general anesthesia and the heterologous blood stored in standard clinical citrate–phosphate–dextrose bags (Baxter) and stored at 4° C for use in EPCC with other pigs within 2 days. The blood type of donors and recipients (i.e., those to be subject to EPCC) was not determined. Lastly, the results reported below were obtained from two further anesthetized female pigs, hereafter referred to as subjects 1 and 2. Their respective weights were 25 and 35 Kg. Of note, in separate work, we found no significant differences in somatic physiological or neurophysiological parameters measured in males and females ^8^. One of these pigs (subject 1) was used to illustrate vascular disconnection at the aorta and the other (subject 2) at the brachiocephalic artery. Because the goal of this work was to arrive at a method for the preservation of native circulation and brain activity, the achievement of this goal was judged from the analysis of the recordings obtained over a sustained period of time (5 hours) from each of these two animals, rather than statistically from a group of animals. Given its exquisite sensitivity to perfusion ^10, 11^, if this duration of neurophysiological activity preservation in the two subjects was achieved under EPCC, the goal was considered as met. This is because the likelihood that our method could faithfully sustain circulation and brain function as measured by chance is negligible. Statistical analyses of several parameters is provided for subjects 1 and 2 to evaluate pre and post EPCC comparisons that informed our goal. Animal housing, environment and nutrition have been previously described ^8^. The following aspects are described in approximate chronological order to facilitate logical progression, but some procedures took place concurrently with or in independence of others where noted.

### 2.3 Monitoring, anesthesia and intravenous fluids

About 30 min prior to surgery, subjects 1 and 2 were sedated with an intramuscular injection of tiletamine and zolazepam (4 - 8 mg/kg of each, in equal amount), atropine (0.04 mg/kg) and buprenorphine (0.05 mg/kg). They were then administered inhaled isoflurane, except during neurophysiological recording as noted below, and oxygen (2 L/ min). These gases were applied first via snout mask and immediately afterwards via endotracheal intubation with mechanical ventilatory support. General anesthesia was maintained throughout the rest of the life of the animals, including euthanasia. This included isoflurane at 1 - 2% v/v with air during non-surgical activities and higher concentrations (up to 5 %) during the performance of incisional or manipulative surgery. The isoflurane was replaced with intravenous anesthesia including ketamine (10-20 mg/Kg/hr), xylazine and guaifenesin using a dose ratio of 1:1:500 during the recording of neurophysiological activity under native and EPCC conditions. The respiratory rate was assisted as needed with a standard animal respirator to maintain 15 - 20 breaths per minute. Intravenous and intra-arterial catheter (auricular and femoral) and esophageal electrocardiogram (EKG) probe and rectal temperature probe insertion was also performed. Heart rate, arterial blood pressure, respiratory frequency, pulse oximetry and capnography were thus monitored to document stability within standard veterinary ranges. An intravenous saline infusion was maintained at 50 mL/ h. During some of the preliminary studies and in subjects 1 and 2, 450 mL of heterologous blood was infused just prior to large artery cannulation. This had no appreciable effect on native blood pressure. This blood had been collected from heterologous pigs and stored at 4°C in citrate–phosphate–dextrose clinical bags (Baxter) for use in EPCC within 2 days.

### 2.4 Cerebral exposure and recording

Each anesthetized pig was first maintained prone on a padded operating table. A strap secured the torso to the table to prevent movement during cranial surgery. The body of the animal was covered with a heated air blanket (Bair hugger, 3M) and the head was surgically draped to allow cranial exposure. Anteroposterior brain length was 5.9 – 6.2 cm for all the pigs weighing 20 - 25 Kg. Thus, the cranium was partially excised as previously, taking into account both this dimension and the provision of electrode coverage over an ample surface of the cerebral convexity ^8^. To this effect, an oval craniectomy, approximately 5 cm rostral-caudal and 4 cm transverse was created. The dura was then opened to allow the apposition of two linear electrocorticography strips to the cerebral surface, each containing 4 electrodes separated by 1 cm, and the near-vertical insertion of two depth electrode linear arrays (Ad-tech), each containing 8 platinum cylindrical recording sites separated by 0.5 cm. These arrays traversed the entire brain in the craniocaudal direction on either side of the midline at the level of the coronal suture. In this configuration, they recorded activity spanning from the immediately subcortical white matter to the hippocampus. Alternatively, in some preliminary study cases the electrodes were placed in the same configuration using 4 burr holes. A Neurovent-PTO 2L brain probe (Raumedic) was inserted into the left frontal lobe at a depth of 1.5 cm for temperature, intracranial pressure, and tissue oxygen saturation monitoring. The scalp was then sutured closed in fluid-tight fashion. The positions of the electrodes and brain probe were documented radiographically aided by a radiopaque ruler. In some preparative cases, subdural radiopaque contrast fluid was also injected to delineate the brain configuration relative to electrode placement. Estimated blood loss for these procedures was 5 mL. Neurophysiological recordings were conducted inside a custom designed Faraday cage using a clinical 32-channel amplifier (Neurofax EEG-1200, Nihon Kohden, Japan). The recording sampling rate was 1000 Hz and a band-pass filter of 0.5 to 300 Hz was applied. All other aspects of these procedures were as described ^8^.

### 2.5 Neurophysiological data analysis

Data from all recording electrodes were analyzed using custom scripts in MATLAB (The MathWorks). This included oscillation frequencies in the range of delta (1 to 4 Hz), theta (5 to 7 Hz), alpha (8 to 12 Hz), beta (13 to 30 Hz) and gamma (31 to 50 Hz) range. Welch’s method for fast Fourier transform was utilized to calculate power for each frequency during 5-min consecutive epochs. To measure temporal changes in frequency spectra, consecutive 10-min epoch depth activity from both pigs was selected and absolute power was calculated for each minute as described ^8^. Parietal strip electrodes were selected for the calculation of power spectral density shown below.

### 2.6 EPCC apparatus

**Figure 1** illustrates the EPCC system. Custom-developed software allowed for multiparameter physiological data visualization in a single interface and in real time via connection to a compactRIO (National Instruments). In this manner, all data including aortic pressure, carotid pressure, pump flow, brachiocephalic artery flow, pump motor velocity, or blood gas and electrolyte measurements were synchronized. The motor part of EPCC was an analog programmable centrifugal pump (BVP-ZX, Harvard Biosciences) that generated pulsatile waveforms. These waveforms were modeled to replicate the native physiological pressure recordings obtained from each individual pig. The BVP-ZX pump controlled perfusion via a magnetic impeller motor that allowed no contact with the perfusate. A commercial pump head with reduced hemolytic action (BP-80, Medtronic) was coupled to the pump. The measured native circulatory pressure waveforms were converted to analog voltage waveforms and used to regulate centrifugal pump speed (RPM). This regulation generated native-like pulsatile flow via custom-developed software using LabVIEW (National Instruments). The correlation between input voltage and RPM was linear. The gains of the input were manually adjusted to ensure adequate pressure and flow while maintaining native-like pulsatile pressure waveforms. Additional custom-developed software allowed the of .CSV files containing a broad range of waveforms derived from native waveforms in order to modify pulsatile systolic and diastolic pressure and beats per minute (BPM) independently of one another or as desired. The custom software also allowed for linear, continuous flow if desired.

**Figure 1.**
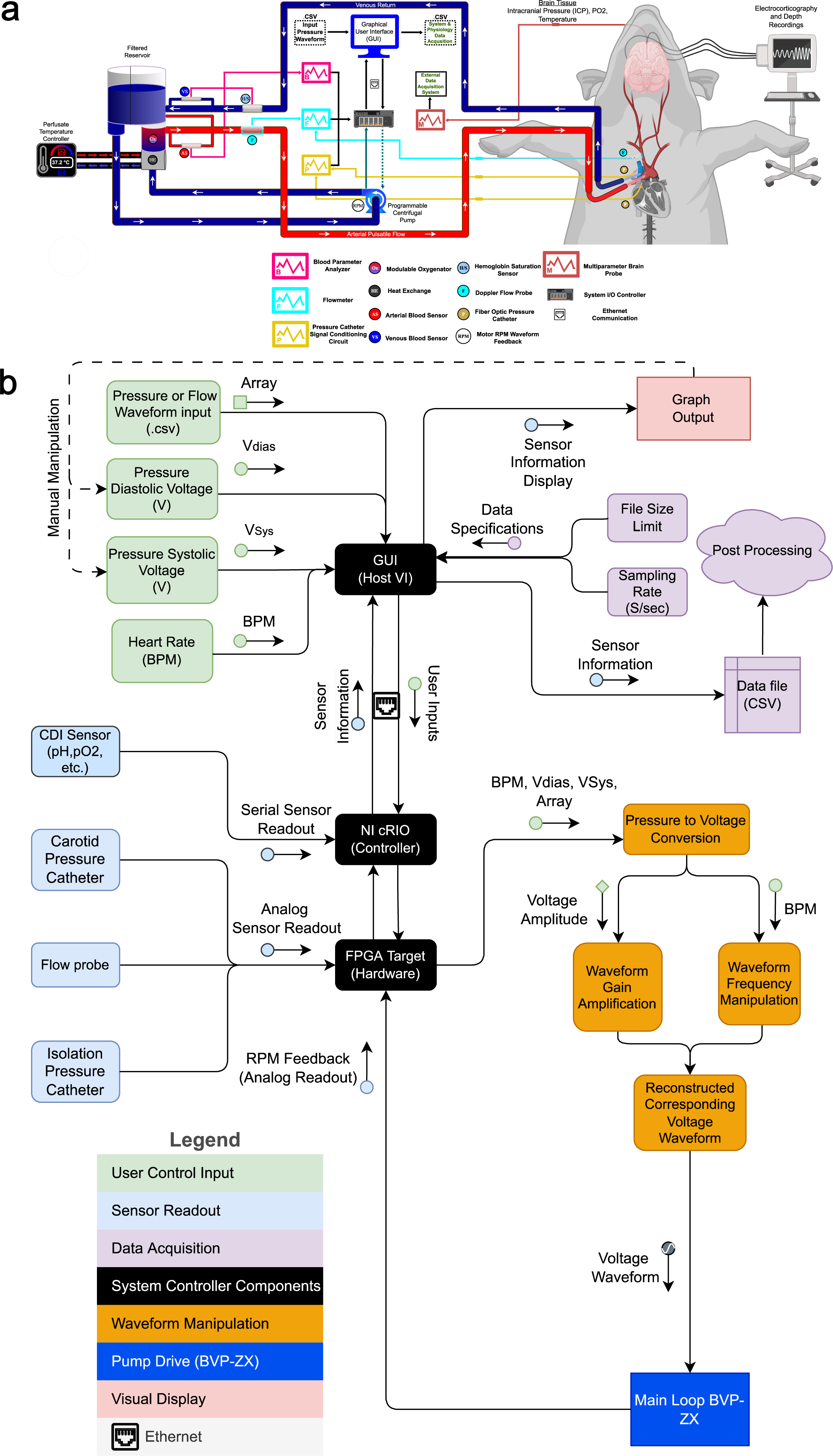
Schematic overview of EPCC and control mechanism. (**a**) Mechanical constitutents of EPCC in relation to blood flow and exchange. Arterial or oxygen-rich and venous or oxygen-depleted blood are represented in red and blue, respectively. (**b**) Representation of the computerized control elements. Relationships between numeric data sources, manipulations and outputs.

The system was complemented with a clinical oxygenator (Affinity NT, Medtronic), an in-line Doppler flowprobe (Transonic), a continuous perivascular Doppler flowprobe (Transonic), a 2 French continuous fiber optic pressure meter catheter (Fiso), a heat exchanger (LT Ecocool), a continuous blood gas monitor (CDI, Terumo) and 3/8 medical grade perfusion tubing (Medtronic). Pump response flow monitoring utilized the inline flow probe located distal to the pump while flow measurement at the level of the brachiocephalic artery used the perivascular flowmeter. The fiber optic pressure monitoring catheter was inserted in-line to the flow of blood. One was placed immediately distal to site of arterial access, with the location is dependent upon surgical approach described below. The second catheter was placed in the right carotid artery just superior to the carotid bifurcation. Venous return from the superior vena cava return was connected to the reservoir to maintain a closed perfusion circuit with the capability to add blood or electrolytes if needed.

Preliminary tests and adjustments of EPCC and its separate constituents were conducted with either water containing polyethylene glycol-200 (MilliporeSigma) dissolved to approximate blood osmolality (300 mOsm ^22^) or pig blood. These studies were performed on the mechanical circuit operating both in isolation and after connection to heparin-treated pig carcasses.

### 2.7 Angiography

Preliminary studies in 3 pigs were used to define the relevant vascular anatomy from the point of cannulation to the small arteries of the brain. This is illustrated in **figure 2** for subject 2, obtained at the end of EPCC. Radiographic contrast agent (Omnipaque, 2% diluted in saline, GE Healthcare) was used in conjunction with an Elite CFD radiographic C-arm (GE Healthcare). At the conclusion of EPCC, a Foley catheter (Bard Medical) was placed inside the brachiocephalic artery and the balloon inflated with air to fill the vessel. An arterial line was then used to inject the contrast agent mixture followed by axial radiography.

**Figure 2.**
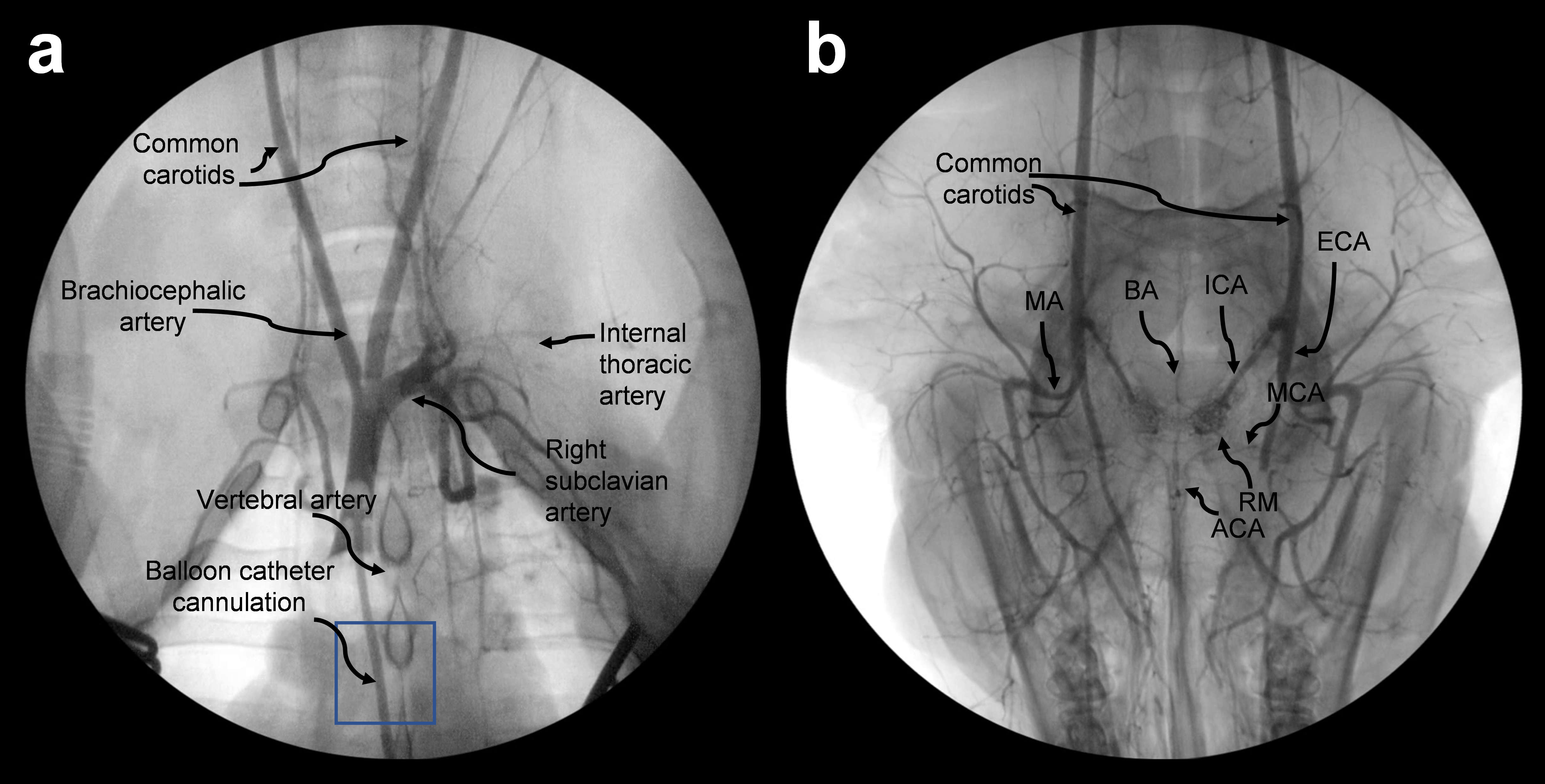
Arterial angiograms of chest and head. Thoracic angiogram obtained via brachiocephalic cannulation (**a**) and extension of the radiographic field to the head (**b**) to illustrate the arteries derived from the carotids. The position of the balloon catheter is indicated in (**a**) with a box. The usual craniocaudal orientation is reversed in (**b**), with the snout pointing toward the inferior part of the image. MA: mandibular artery; BA: basilar artery; ICA: internal carotid artery; ECA: external carotid; MCA: middle cerebral artery; ACA: anterior cerebral artery; RM: rete mirabile.

### 2.8 Carotid artery recordings

Carotid artery exposure and cardiovascular surgery followed supine inversion and positional securing of the pigs as above. The snout was maintained at heart level using a cushioned support. Of note, while the human brachiocephalic, left common carotid, and left subclavian arteries emerge separately from the ascending aorta, pigs possess only two branches. The brachiocephalic artery gives rise to not only the right subclavian and right common carotid arteries, but the left common carotid artery as well ^23^. Thus, from a cerebral perfusion perspective, access to the brachiocephalic artery is important to ensure viability.

The right common carotid artery was dissected using electrocautery. Following exposure of 7 - 9 cm of the artery, the fiber optic catheter was transiently threaded through a 22 G arterial puncture cannula and left in place 1 - 2 cm deep, with the tip pointing towards the brain, thus allowing the continuous measurement of arterial pressure before and during EPCC. Data acquisition was immediately initiated through interfacing the corresponding signal conditioning circuit with the LabVIEW controlled Compact-RIO 9045 for direct visual analysis and data storage. This allowed the use of carotid native recordings in subsequent comparisons with EPCC recordings. Only as a backup to the main pressure data acquisition system was proprietary Fiso software concurrently used.

### 2.9 Native thoracic artery recordings

Simultaneously with or following carotid artery access, the thoracic cavity was exposed via a median sternotomy. The pericardium was opened with electrocautery and the heart and great vessels were exposed. Afterwards, a second 2 French fiber optic pressure catheter was similarly inserted into either the ascending aorta (subject 1) or the brachiocephalic artery (subject 2) for continuous native pressure measurements. The perivascular Doppler flowprobe was fitted to encircle the brachiocephalic artery and immersed in a small amount of surgilube ultrasound gel (HR Pharmaceuticals). Continuous native pressure and flow recordings (when applicable) were visually displayed and stored through the Compact-RIO 9045 interface. Since circulatory stability was documented in preliminary studies lasting over 4 hr, both probes were removed following 5-10 min of data collection. A single representative native pressure waveform and the pulsation frequency were isolated using MATLAB (MathWorks) and then used as input to the pulsatile pump as a .CSV file. The subjects were then systemically anticoagulated with 300-400 U/kg of heparin prior to EPCC initiation.

### 2.10 EPCC circuit priming

The circuit was primed separately from but simultaneously with the performance of vascular access surgery to minimize blood loss. Prior to the addition of heterologous blood, the Affinity NT reservoir was primed with 1000cc of 0.9% Sodium Chloride solution. 5,000 U of heparin were added. Pump-induced positive pressure established flow from the reservoir to the heat exchanger/oxygenator and then to the rest of the vascular system upon connection. Priming ensured proper fluid-solid interfacing with the tubing devoid of air bubbles. Just prior to EPCC, 600 mL of heterologous blood was deposited into the venous reservoir. The blood was filtered and continuously recirculated in a closed loop fashion prior to initiation of EPCC to remove air bubbles that may develop during reservoir decantation. Following circuit operation for up to 30 min prior to EPCC, the passing of blood through the heat exchanger/oxygenator ensured an initial and stable temperature of 37.2 °C and a fractional oxygen percentage of 80% was supplied at 1.5 liters per minute of oxygen flow across the oxygenator. The delivered oxygen percentage and sweep gas flows were then adjusted hourly based on blood gas measurements to maintain physiologic pH and oxygen levels that ensured fully saturated hemoglobin. Arterial cannulation was then performed in the distal ascending aorta or proximal arch with an aortic size matched cannula to allow innominate artery or arch isolation for subsequent EPCC initiation. Venous cannulation was performed through the right atrial appendage with an appropriately sized dual stage venous cannula (Medtronic). Cardiopulmonary bypass was then commenced prior to transitioning to EPCC.

### 2.11 Aortic isolation (subject 1)

**Figure 3** illustrates the connection of the aorta following its isolation. The ascending aorta was cannulated with an 8 French fenestrated arterial cannula (Medtronic) approximately 5 cm distal to the aortic valve and coronary arteries and secured in place. A clamp was applied across the proximal ascending aorta and proximal descending thoracic aorta as depicted in Figure 3 a. This prevented retrograde flow, thus ensuring devitalization of the heart. The subclavian arteries were not ligated to ensure preservation of the vertebral arteries. A venous straight cannula (Medtronic) was placed in the superior vena cava and secured and snared to accept only flow returning from the superior vena cava. As noted, the aortic clamp also occluded the proximal descending thoracic aorta immediately past the origin of the left subclavian artery, eliminating perfusion to the descending aorta. Lastly, to prevent flow to the upper extremities, both subclavian arteries were occluded distal to the origin of their respective vertebral artery. The 2 French thoracic pressure catheter was then reintroduced to initiate isolated aortic recordings. Estimated blood loss for this and the following procedure was 50 mL. The connections thus created between arterial and venous cannulas and the cardiopulmonary bypass tubing allowed for full EPCC.

**Figure 3.**
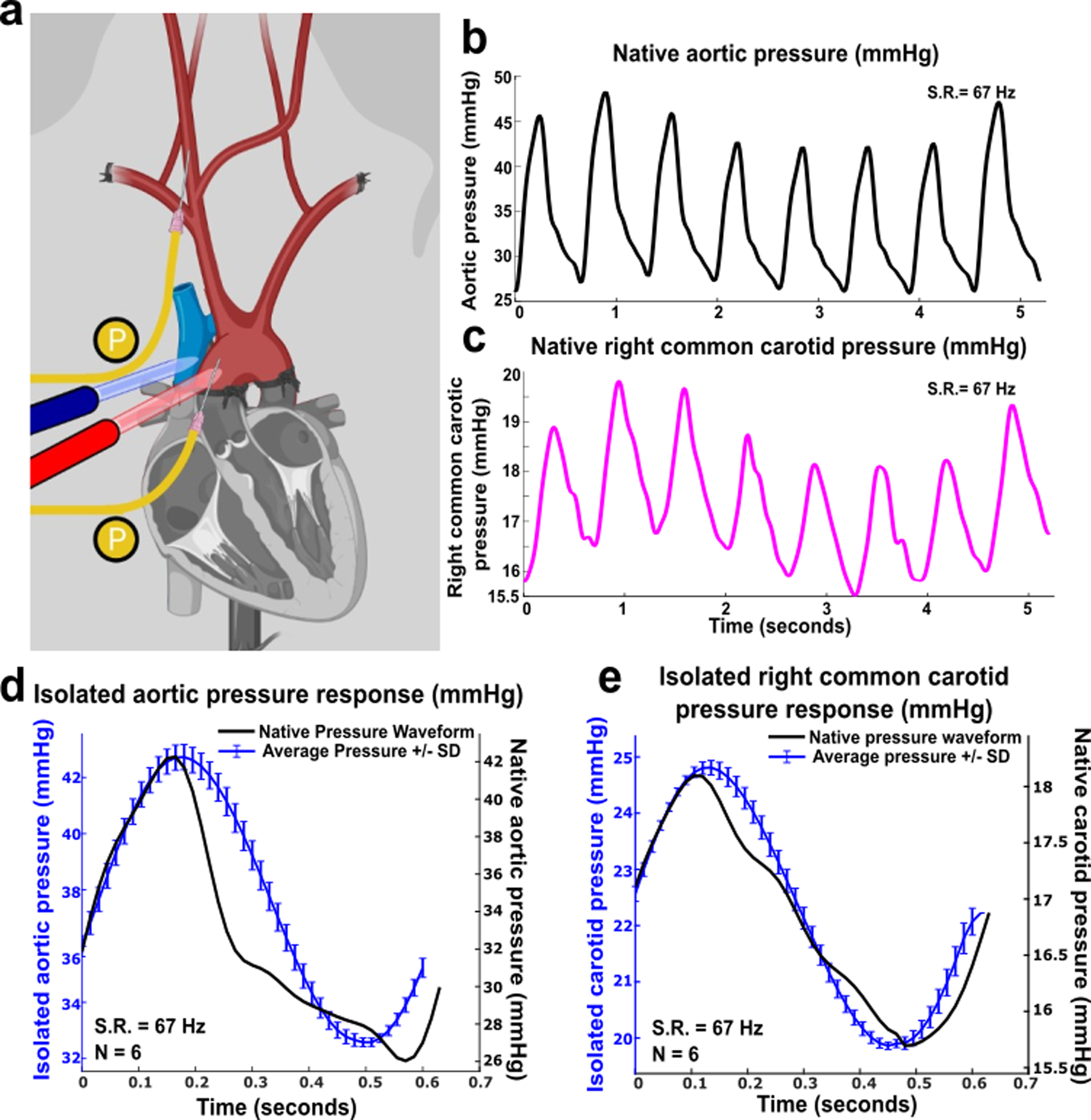
EPCC pressure relative to native perfusion pressure following aortic isolation. (subject 1). (**a**) Vascular structure and aortic isolation. Red: arteries; blue: veins. P (yellow) indicates pressure measurement locations. (**b**) Native aortic pressure. (**c**) Native right common carotid pressure. (**d**)-(**e**) Comparative waveforms of 6 averaged recordings under EPCC (designated here as isolated condition) with native waveforms used as input for the aortic pressure and for the right common carotid pressure, respectively. All recordings were acquired at 67 samples per s. S.R.: sampling rate. In (**d**)-(**e**) the left vertical axis is colored in blue, as is the arterial pressure measured under EPCC.

### 2.12 Brachiocephalic artery isolation (subject 2)

Figure 4 shows the connection established after brachiocephalic isolation. The procedure was similar to the aortic isolation except for the placement of the arterial cannula, pressure catheter and vascular clamps and incorporation of the perivascular flow probe. In this animal, the aorta was cannulated with an 8 French fenestrated arterial cannula in the transverse arch near the origin of the brachiocephalic trunk. A venous straight cannula was secured in the superior vena cava and snared. The ascending aorta was clamped proximal to the origin of the brachiocephalic trunk and also clamped in the transverse arch between the brachiocephalic trunk and left subclavian artery, preventing flow to the left subclavian artery and lower extremities. Both subclavian arteries were occluded distal to the origin of the vertebral arteries. Of note, whereas the right and left vertebral arteries join to form the basilar artery, the left cerebellar hemisphere might experience altered perfusion, although circle of Willis anastomoses may compensate for this phenomenon. In contrast with the aortic isolation, the 2 French pressure catheter was placed immediately proximal to the flow probe at the base of the brachiocephalic branch.

**Figure 4.**
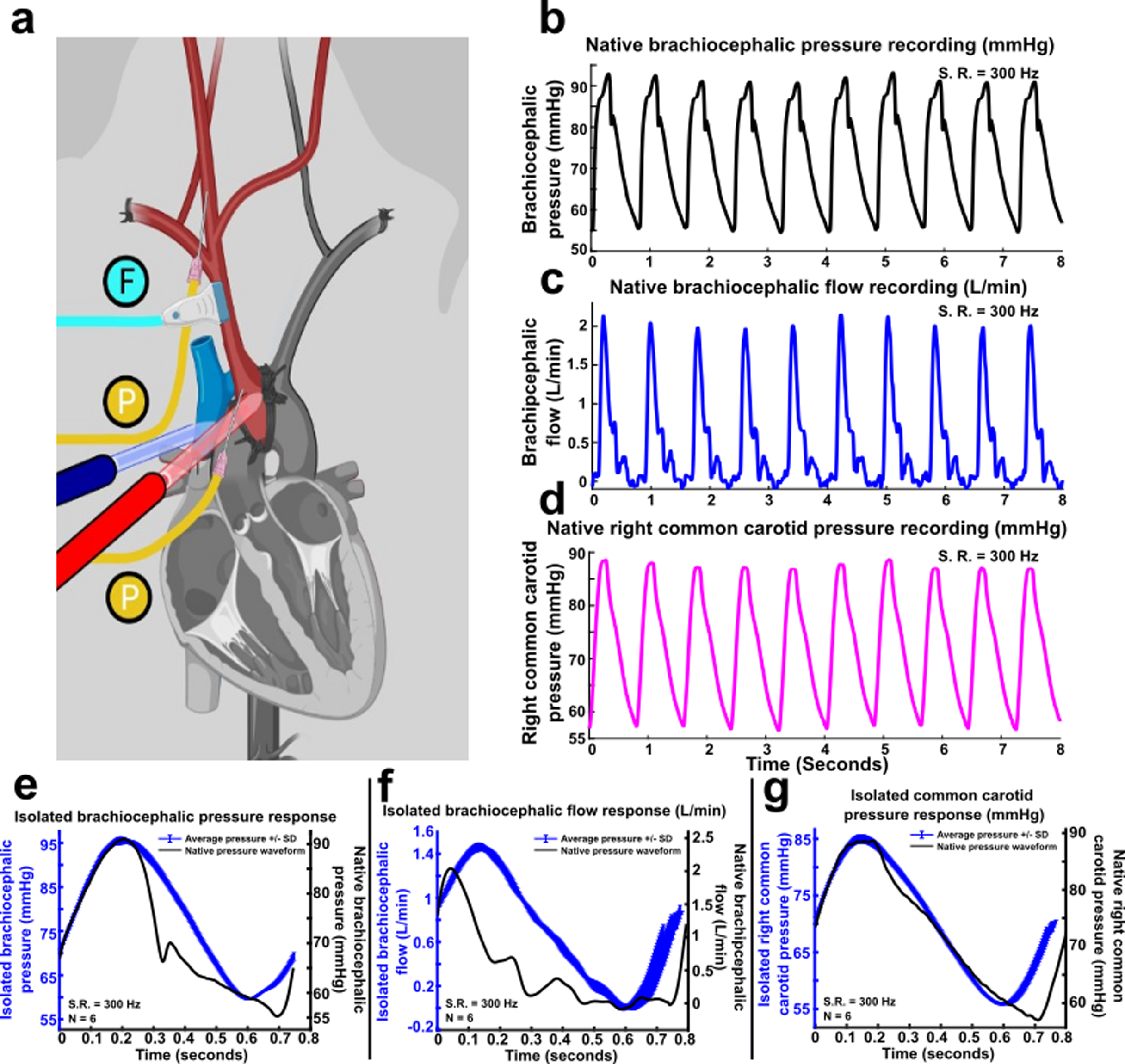
EPCC pressure relative to native perfusion pressure following brachiocephalic aortic isolation. (subject 2). (**a**) Vascular structure and brachiocephalic isolation. Red: arteries; blue: veins. P (yellow) indicates pressure measurement locations. (**b**) Native brachiocephalic pressure. (**c**) Native Doppler-measured brachiocephalic flow. (**d**) Native right common carotid pressure recordings. (**e**)-(**g**) Comparative waveform analysis of 6 average recordings under EPCC (designated here as isolated condition) and sampled native waveforms used as input for the brachiocephalic pressure, brachiocephalic flow and right common carotid pressure, respectively. All recordings were sampled at 300 samples per s. S.R.: sampling rate. In (**e**)-(**g**) the left vertical axis is colored in blue, as is the arterial pressure measured under EPCC.

### 2.13 EPCC initiation and data acquisition

Following initial stable cardiopulmonary bypass initiation, pulsatile flow was initiated and pressure readings as detailed below were obtained. As noted above, the native pressure waveform and BPM were used for input into the custom LabVIEW program to initiate EPCC. The carotid pressure, aortic or brachiocephalic pressure, flow, and pump motor performance were collected through the LabVIEW interface and stored as a .CSV file for subsequent analysis. Any overt blood loss was measured and compensated via supplementation of heterologous blood using the venous reservoir. Maximum blood replenishment was 1.6 L in the course of 4 hr.

Maximum EPCC duration was 5 hr. The following principal parameters were measured. Subject 1: Carotid pressure (mm Hg), aortic pressure (mm Hg), and motor response (RPM), sampled at 67 Hz, and blood gas and electrolyte analysis at 1 sample/min. Subject 2: Carotid pressure (mmHg), aortic pressure (mmHg), brachiocephalic flow, and motor response (RPM), sampled at 300 Hz, and blood gas and electrolytes as above.

### 2.14 Additional hematocrit, gas and electrolyte analyses

**Supplementary tables 1** and **2 online** illustrate additional blood composition analyses. To assess the consistency of continuous CDI blood monitoring, blood samples were collected from the reservoir for analysis every 15 min. This included portable clinical analyses (sodium, potassium, chloride, CO_2_, anion gap, ionized calcium, glucose, urea nitrogen, creatinine, hematocrit and hemoglobin; I-stat 1, Abbott) and separate conventional clinical laboratory analyses of complete blood count and blood chemistry obtained after EPCC.

### 2.15 Termination

Euthanasia was induced at the conclusion of each study under general anesthesia by the intravenous addition of pentobarbital at excess dose (120 mg) sufficient to produce asystole, cessation of spontaneous respiration and the development of fixed and dilated pupils with absent corneal reflexes. Necropsy was performed to verify electrode placement and brain configuration and integrity.

### 2.16 Histological examination of the brain

Upon euthanasia, the brains were subject to measurement of cerebral cortex thickness and layer organization following standard formalin fixation and paraffin embedding. Coronal sections were exposed to Nissl/ luxol fast blue (Nissl/LFB) as well as luxol fast blue/ periodic acid Schiff (LFB/PAS) stains according to published methods ^24, 25^. In detail, serial 5-µm sections were prepared by rotary microtome, verified by for quality under dark-field microscopy ^26^ and air-dried before being deparaffinized and run to 95% ethanol. Sections for both stains were impregnated overnight in a common vessel with 0.1% w/v solution of luxol fast blue prepared in 0.05% acetic acid and 95% ethanol at 58°C. Slides for both stains were briefly destained in 95% ethanol, and rinsed in dH_2_O the next morning, before differentiation in 0.05% lithium carbonate followed by further differentiation in exchanges of 70% ethanol. Slides for the respective combo-stains were then placed in separate containers of dH_2_O for separate, but same day finishing. Nissl/LFB slides were decanted free of dH_2_O and were subjected to hot, acidified 0.1%Cresyl-Echt-Violet stain for 10 minutes, before dH_2_O rinse, air-drying, and rapid, stain-preserving-movement through alcohols and xylenes, and coverslipping with synthetic mounting media. LFB/PAS slides wre decanted free of dH_2_O and oxidized in 0.5% periodic acid for 10 minutes, rinsed in dH_2_O, and reacted with commercial Schiff’s reagent (Sigma-Aldrich) for 15 minutes. Following Schiff’s-aldehyde formation, slides were rinsed in dH_2_O and nuclei were counterstained with commercial hematoxylin (Selectech 560, Leica Microsystems). Following hematoxylin, slides were blued in running tap water for 5 minutes, then moved through alcohols and xylenes, before coverslipping with synthetic mounting media. Cortical layers were examined and compared with the layering of the human cortex as described ^12^. The sections were photographed on a Leica DM2000 microscope equipped with a Jenoptik Gryphax NAOS CMOS camera.

## 3. Results

### 3.1 Surgical approach

Cranial surgery was uneventful in all preliminary study pigs and in subjects 1 and 2. One preliminarily studied pig harbored a 3 cm right cerebral cortex tumor and one was subclinically epileptic (i.e., no seizures had been previously apparent) as noted by electrocorticography. There were no appreciable untoward effects associated with the use of heterologous blood. Consistently with the known propensity to arrythmias with cardiac manipulation of some domestic pigs ^27^, one preliminary study pig experienced ventricular fibrillation upon sternotomy and was terminated.

### 3.2 Perfusate composition stability

One preliminarily studied pig received 100 mEq of sodium bicarbonate into the blood reservoir to rectify a clinically significant reduction in blood pH (from 7.2; base deficit −5 mmol/L). All the blood samples analyzed in a clinical veterinary laboratory for subjects 1 and 2 yielded values within acceptable clinical veterinary ranges of variation relative to native samples. **Supplementary tables 1** and **2** report these values. Serial blood samples studied at the bedside (I-stat analyzer, **supplementary table 3**) also illustrated negligible or clinically acceptable variations in blood pH, CO_2_, O_2_, sodium, potassium, glucose and other analytes. This also suggests that the blood supplemented into the reservoir was sufficient to maintain the metabolic homeostasis of the perfused tissue.

### 3.2 Pressure and flow at the carotid arteries

#### 3.2.1 Subject 1

Native aortic pressure recordings measured from subject 1 for the purpose of physiological replication under isolated conditions were sampled at 67 Hz. An 8-iteration (i.e., 8-beat) recording is depicted in **figure 3B**. Under native conditions, subject 1 exhibited an average systolic pressure of 44.5 ± 2.5 (mean ± SD) and an average diastolic pressure of 26.7 ± 0.7 mmHg. Native average mean arterial pressure (MAP) was 32.7 ± 1.3 mmHg, with a consistent heart rate of 98.7 BPM. As stated above, these pressures were below those typically measured in other animals but we did not modify them toward greater values since, as shown below, there was no appreciable impact on neurophysiological activity. In fact, this pressure context allowed us to pursue the goals of this study, i.e., reproducibility of any native pressure values and preservation of neurophysiological activity, under these lower values. A representative waveform with a systolic and diastolic pressure of 42.5 and 26.1 mmHg, respectively, was used as an input waveform for pump control. The MAP was calculated at 31.6 mmHg. The response to the pump input measured in the aorta is depicted in **figure 3D** at the specified heart rate of 98.7 BPM. 6 consecutive waveforms were averaged for further analysis. This indicated that the aortic pressure under EPCC was, on average, 42.7 mmHg for the systolic and 32.6 for the diastolic level, yielding a MAP of 36.0 mmHg. RPM analysis indicated that the pump consistently provided cycles between 1754.2 and 1359.6 revolutions per minute to generate these systolic and diastolic pressures.

Carotid pressure recordings were also sampled at the same rate. **Figure 3C** represents a corresponding 8-iteration of native carotid pressure recordings in response to the corresponding aortic pressure fluctuations noted. Significant dampening of the carotid pressure waveform was observed with an average systolic pressure of 18.9 ± 0.7 mmHg and average diastolic pressure of 16.2 ± 0.5 mmHg. Average MAP was 17.1 ± 0.5 mmHg. Measurements were taken in the right common carotid artery at the site of native recordings. The systolic and diastolic carotid pressure observed corresponding to the aortic waveform input waveform aforementioned was 18.2 mmHg and 16.0 mmHg, respectively, providing a MAP of 16.7 mmHg. The carotid pressure response under EPCC in comparison with the native pressure waveform is illustrated in figure 3E. 6 consecutive waveforms were averaged for further analysis corresponding to the aortic recordings aforementioned. This indicated that the EPCC pressure response in the carotid artery resulted in a systolic/diastolic ratio of 24.8/ 19.9 mmHg, thereby resulting in a MAP of 21.51 mmHg.

#### 3.2.2 Subject 2

Native and EPCC brachiocephalic and carotid pressure recordings obtained from subject 2 were recorded at a sampling rate of 300 Hz. A subset of native representative physiological recordings (10 iterations) taken for EPCC replication are depicted in **figure 4B-D**. Brachiocephalic pressure, measured near the origin of the artery, consisted of an average systolic to minimum diastolic pressure ratio of 91.7 ± 0.93 to 55.1 ± 0.68 mmHg, with an average MAP of 67.3 ± 0.63 mmHg. Native heart rates were recorded as consistently being between 75 and 76 BPM. Native brachiocephalic flow was 6.42 mL per beat. Right common carotid native recordings exhibited undiminished tonicity, providing a uniform average systolic to diastolic pressure ratio of 87.5 ± 0.66 to 57.0 ± 0.54 mmHg, with an average MAP of 67.2 ± 0.46 mmHg, similar to the values obtained in the brachiocephalic recordings.

**Figure 4E-G** represents a waveform comparative analysis between native and EPCC pressure and flow to illustrate mechanical fidelity. As aforementioned, EPCC using a preset native waveform recorded from subject 1 was amplified to allow for characteristic native systolic/ diastolic pressure and flow ratios. An approximate number of 80 BPM was selected for validation. Minimal fluctuations from native readings were noted as demonstrated in each respective comparative analysis. The results indicate that, under EPCC, using an input pressure waveform with an adjusted systolic/diastolic ratio of 91.0/ 55.3 mmHg and a MAP of 67.2 mmHg corresponding to native recordings, the output brachiocephalic pressure recordings (evaluated using 6 averaged waveforms) had a systolic to diastolic pressure ratio of 95.5/ 59.6 mmHg with a MAP of 71.6 mmHg. In response to EPCC brachiocephalic waveform input with an expected corresponding native carotid pressure of 88.7/ 57.0 mmHg and a MAP of 67.6 mmHg, the EPCC carotid pressure was, on average, 85.0/ 55.9 mmHg with a MAP of 65.6 mmHg. Flow analysis using 6 averaged pressure waveforms yielded 8.49 mL per beat. The maintenance of the desired brachiocephalic and common carotid pressures was enabled by a consistent RPM systolic-diastolic ratio of 2535.7/ 1463.0 RPM.

While the significant pressure waveform similarities noted between native and EPCC conditions indicate that both aortic and brachiocephalic flow approaches allow for the faithful replication of native waveforms, the results indicate that control of flow through the brachiocephalic approach leads to limited pressure dampening between the arterial cannulation site and the common carotid artery. Additionally, the ability to maintain almost identical systolic/ diastolic pressure to native conditions as a result of direct vascular access under EPCC suggests the superiority of the brachiocephalic approach.

### 3.3 Cerebral activity under EPCC

EPCC was compatible with the sustained preservation, without interruption, of electrical activity both in subject 1 (**figure 5**) and subject 2 (**figure 6**). **Figures 5A** and **6A** compare recording traces before and after EPCC, illustrating the similarity in amplitude and frequency. The quantification of the resulting absolute power spectra per frequency before and after EPCC is shown in **figures 5B** and **6B**. To illustrate this further, we calculated power spectra of one minute depth activity epoch for each EEG frequency, which included delta (1-4 Hz), theta (5-7 Hz), alpha (8-12 Hz), beta (13-30 Hz) and gamma (31-50 Hz) frequencies. Post EPCC, there was little to no change in absolute power except for noise apparent in subject 1, which was later suppressed by regrounding the electronic equipment (**figure 5C** and **6C**). In both cases, and despite differences in native blood pressure, this activity was normal ^8^. Both craniotomy and burr hole approaches allowed for virtually indistinguishable recordings. However, as expected, the latter approach allowed for the preservation of a greater intracranial pressure by about 5 mmHg, which may be relevant to maintain cerebral perfusion pressure commensurate with the likely native pressure. Of note, characteristic electrocorticography and depth recordings from the pig brain were previously illustrated using craniotomy and other recording means essentially identical to the present study and this allowed comparison of those previous recordings with human awake electrocorticography. Despite the normal degree of individual variability inherent to absolute neurophysiological signal power analysis, the results from subjects 1 and 2 and from three additional pigs preliminarily studied for shorter times were not appreciably different from the previous human and pig recordings. This comparison is not shown since the publication is available ^8^. Thus, we could not discern any neurophysiological impact of EPCC.

**Figure 5.**
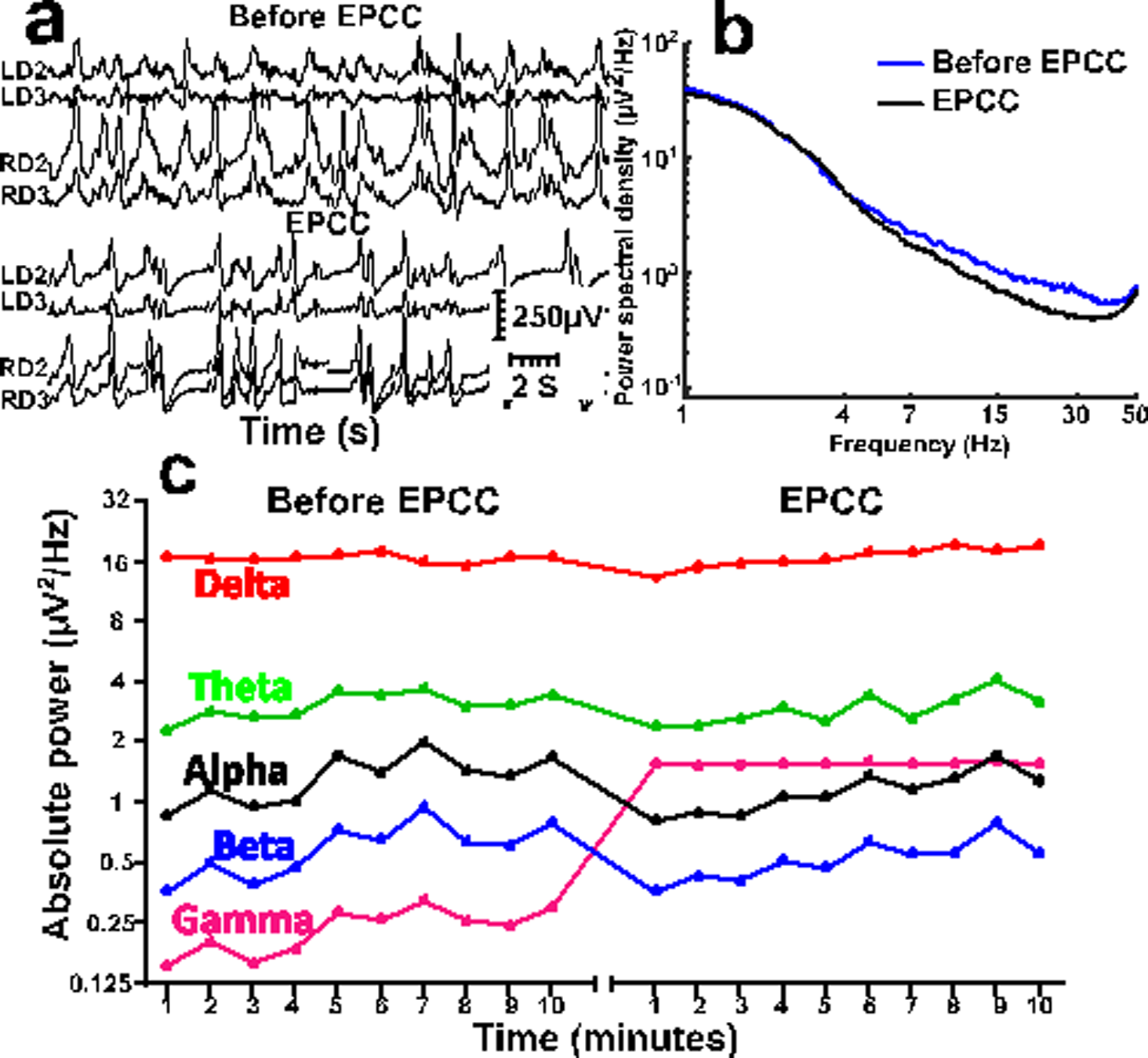
Comparison of depth neurophysiological activity before and after aortic EPCC. Depth recordings from subject 1 before and after EPCC. (**a**) Depth activity from left (L) and right (R) depth (D) electrodes corresponding to subcortical brain regions spanning from the subcortical white matter (LD2 and RD2) to the dorsal striatum (LD3 and RD3) before and following 4 hours of EPCC. (**b**) Absolute power spectral density of native (blue spectra), post-EPCC (black dotted spectra) LD2 depth recordings in subject 1. (**c**) Absolute power spectra of delta (red), theta (green), alpha (black), beta (blue), and gamma (magenta) activity defined as in standard electroencephalography and measured in LD2 using 10-minute epochs under native perfusion and under EPCC. Electrical noise (50 Hz) interfered with the accurate measurement of gamma frequency after EPCC.

**Figure 6.**
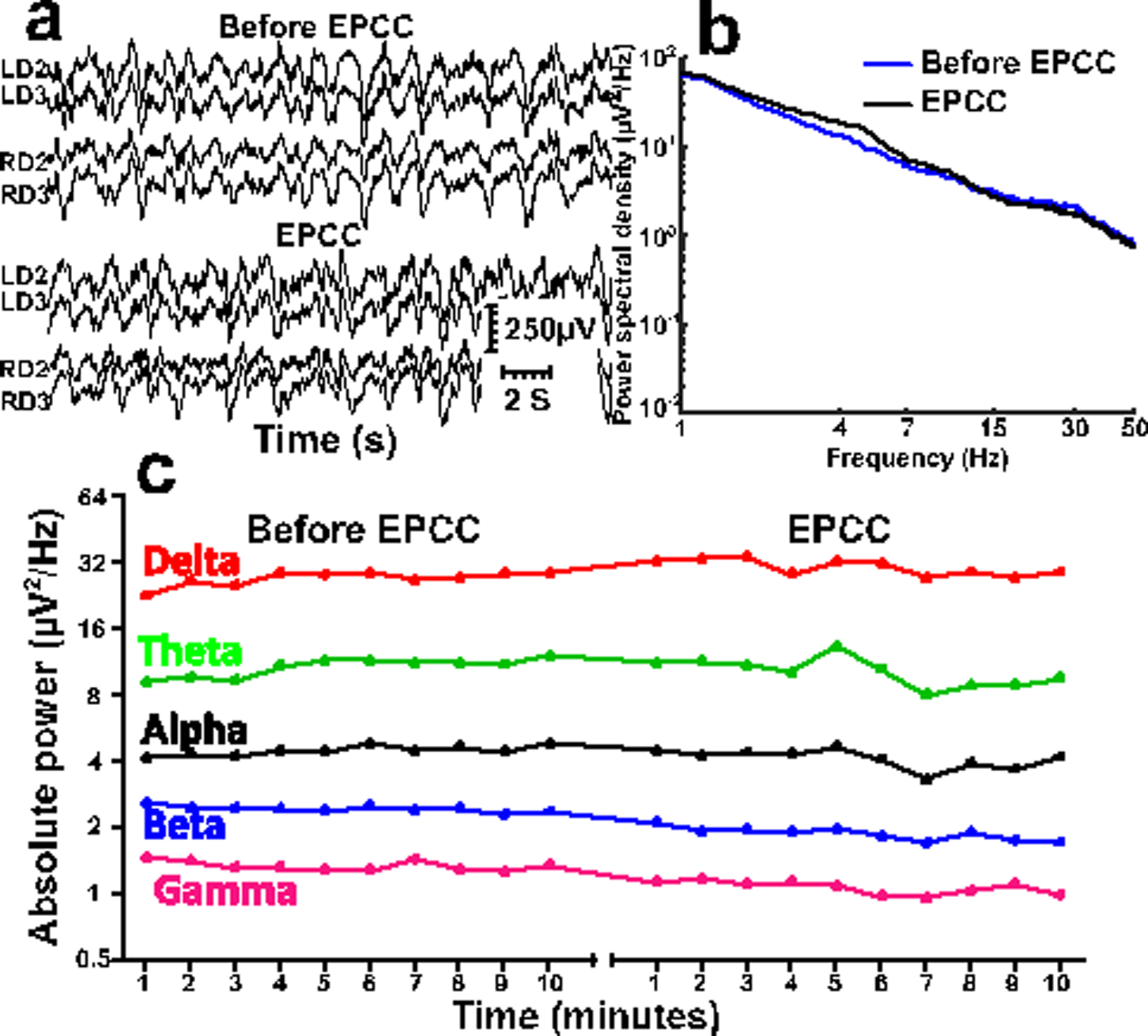
Comparison of depth neurophysiological activity before and after brachiocephalic EPCC. Depth recordings from subject 2 before and following 4 hours of EPCC. (**a**) Depth activity from left (L) and right (R) depth (D) electrodes corresponding to subcortical brain regions spanning from the subcortical white matter (LD2 and RD2) to the dorsal striatum (LD3 and RD3) before and following EPCC. (**b**) Absolute power spectral density of native (blue spectra), post-EPCC (black dotted spectra) LD2 depth recordings in subject 1. (**c**) Absolute power spectra of delta (red), theta (green), alpha (black), beta (blue), and gamma (magenta) activity defined as in standard electroencephalography and measured in LD2 using 10-minute epochs under native perfusion and under EPCC.

Physiologic intracerebral pressure and temperature were also maintained. Tissue oxygenation, however, was greater than in native conditions, likely because of supplemental oxygen administration (**figure 7**).

**Figure 7.**
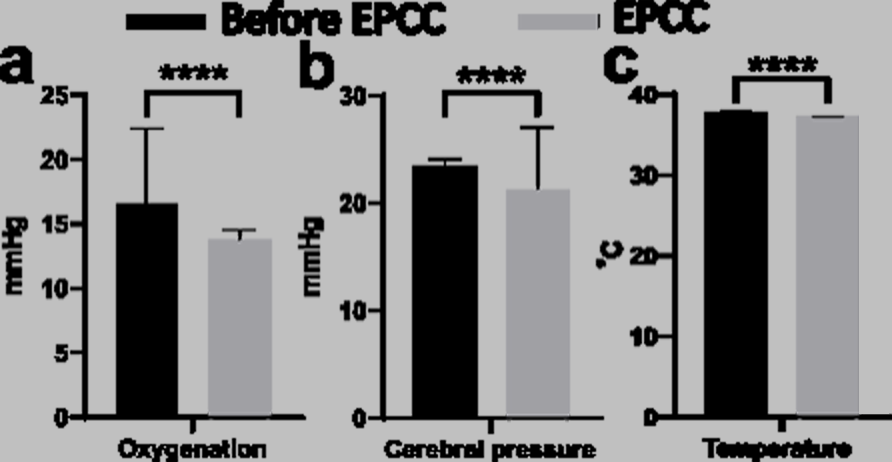
Physical characteristics of cerebral tissue after EPCC. Brain probe measurements of frontal lobe cerebral oxygenation (**a**) (mmHg of oxygen), barometric pressure (**b**) (mmHg) and temperature (**c**) (°C) in subject 2. Black bars represent average and SD of measurements obtained in the native, pre-EPCC state for 10 minutes with a sampling rate of 1 Hz. Gray bars indicate averaged values of 10 minutes epochs under EPCC, measured over 5 hr. **** represents a significance of p<0.0001, unpaired *t*-test.

### 3.4 Microscopic structure of the brain

Consistent with previous studies of brain relative to body weight ^28^, brain weights were 60 g and 63 g for subjects 1 and 2, respectively. Sections from several brain regions demonstrated preservation of cellular elements after EPCC relative to untreated pigs. Cortical cell structure and layer distribution (exemplified in **figure 8** for the somatosensory cortex, with additional normative examples and human comparison in ^8^) was unaltered by light microscopy histopathologic evaluation using both Nissl/ luxol fast blue and luxol fast blue / periodic acid Schiff staining. Specifically, there were no abnormalities of either the gray or white matter.

**Figure 8.**
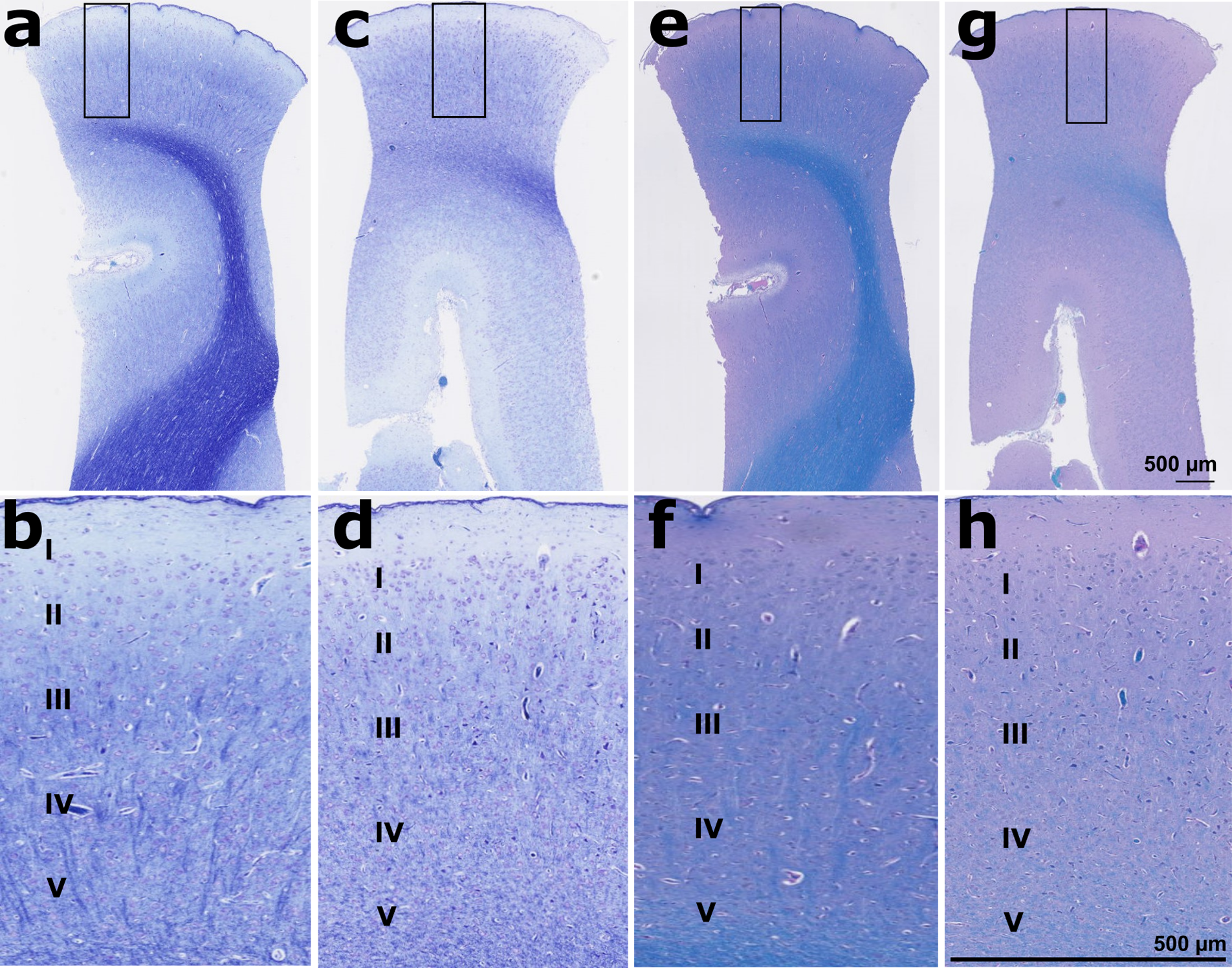
Microscopic structure of the cerebral cortex after EPCC. A and B: Nissl / luxol fast blue-stained cortical structure of subject 1 after EPCC. C and D: Nissl / luxol fast blue-stained cortical structure of subject 2 after EPCC. E and F: luxol fast blue / periodic acid Schiff-stained cortex of subject 1. G and H. luxol fast blue / periodic acid Schiff-stained cortex of subject 2. Section thickness: 5 µm, photographs obtained with 2.5 × and 10 × objectives.

## Discussion

Previous primate and canine studies attempted to preserve brain viability after isolation of the cerebral circulation. No neurophysiological measurements were reported pre-circulatory intervention in the canine model, thus limiting post-perfusion inferences ^5, 6^. The monkey studies documented a limited repertoire of non-quantified neurophysiological activity ^3, 4^. Thus, no direct comparisons with our study are feasible. Another difference between these studies and our approach is our use of an elevated sampling rate for all electrocorticography and deep brain structure recordings coupled with robust noise reduction techniques that allow for the measurement of signals particularly vulnerable to metabolic or other forms of neural dysfunction ^8, 29–31^.

The preservation of cerebral activity as studied by our methods under EPCC was maintained over the duration of each subject study (5 hours). Except for excess brain tissue oxygenation upon oxygen supplementation, and mild intracranial pressure changes when craniotomy was used, this system was associated with near-native levels of cerebral physiological parameters such as intracranial pressure, tissue oxygen saturation and temperature.

We measured the neurophysiological activity recordable by contact with the cerebral cortex and with deep structures particularly susceptible to ischemia ^10, 13^. The division of this activity into standard spectral oscillatory frequencies and the selectivity and velocity of changes noted for some of these frequencies after ischemia allowed us to conclude that EPCC imposes no significant circulatory changes on the brain relative to native perfusion. In fact, because even profound neurophysiological changes during carotid endarterectomy can prove fully reversible ^10^, no cellular injury is expected from EPCC. Thus, our approach merits consideration for the investigation of cellular signaling and other mechanisms likely to be perturbed under linear flow conditions. Of note, in the pig, and in contrast with other animals, carotid artery flow is closely correlated with cerebral blood flow ^32^. As such, our measurements at the carotid artery are likely to directly impact cerebral blood flow.

EPCC is advantageous compared to uniform or continuous perfusion from a neurological perspective. Experience with cardiopulmonary bypass and extracorporeal membrane oxygenation suggests that continuous flow results in impairment of cerebral autoregulation and this may limit certain studies ^33^. Left ventricular assist device patients subject to continuous flow mode also exhibit differences in cerebral blood flow velocities and increased sympathetic activity, although autoregulation may be preserved ^34^.

Since we did not measure blood flow to the head without the brain, we can only estimate the distribution of perfusion to the brain relative to the head under EPCC. Yet, such an estimate offers context for future work. Although we did not determine the anatomical demarcation of the perfused relative to the devitalized parts of the head, we estimate from the preliminary studies that, for the pig weights that we studied, the weight of the entire head does not exceed 1.5 Kg. Similarly, brain weight is near 60 g. Thus, assuming a cerebral blood flow of approximately 90 mL/ min per 100 g ^32, 35^, and a head (without the brain) blood flow of 10 mL/ min per 100 g ^35^, we estimate an EPCC blood flow of 150 mL/ min to the head (except the brain) and 54 mL/ min to the brain. These estimates depend on the reliability of blood flow measurements. For example, human cerebral blood flow, which has been more extensively studied than pig cerebral blood flow, has long been estimated at only 50 mL/ min per 100 g ^36, 37^. Further, the estimates are also subject to significant pig breed-specific head tissue composition (which primarily includes bone and muscle) variability. However, the relative tissue abundance of the head can be determined via several methods. Moreover, blood flow to these individual tissues has been estimated for the pig ^35^. Therefore, it is possible, after these calculations, to greatly simplify metabolic tracer or pharmacological kinetic studies using EPCC. Further, the blood distribution volumes and metabolic activities of bone and resting head muscle are far below those of other organs such as liver and kidney, thus reducing the additional physiological complexities that curtail such studies.

Upon artificial perfusion, isolated pig brains can retain both cellular configuration and activity for several hours post-mortem ^7^. However, the electrocorticogram remains isoelectric. In these studies, the relative contribution to this isoelectricity of the perfusate composition and of the intrinsic suppression of field potentials after artificial perfusion was not quantified. However, it is likely that the electrocorticogram, which reflects the summation of a large ensemble of field potentials not made apparent by single-unit and other localized recordings, indicates cessation of synaptic critical for high-order network activity. This is implicit in the use of the loss of the electroencephalographic signal for the diagnosis of brain death.

### Potential limitations of the study

We did not study EPCC for durations greater than 5 hours although our evidence suggests that longer artificial perfusion times are feasible. Correction of blood electrolytes or hematocrit was not necessary in this study interval other than bicarbonate in one case, likely because of heterologous blood replenishment of blood losses. We assume that maintenance for prolonged times of infra-physiological cerebral perfusion pressure due to craniotomy may result in cerebral edema. In our study, no edema was appreciated by histological analysis. Further, because the brain is exposed and directly visible, and also under direct and operative microscope visualization for the procedure, we could have detected any significant edema based on our comparative human neurosurgical expertise. Lastly, EEG is also rapidly sensitive to early cerebral edema induced by brief perfusion failure, and this occurs in correlation with histological findings ^38^.

We used ketamine anesthesia during the acquisition of the neurophysiological recording segments used for comparison with previously established data ^8^. Ketamine was preferred over other anesthetics given its reduced impact on brain activity, including a lack of depression of the cerebral oxygen metabolic rate. Rather, ketamine increases oxygen consumption slightly in association with increased metabolite supply ^39^. This may amount to a ∼15% increase in regional glucose metabolic rate (when studied at sub-anesthesia concentrations) but altered coupling or disequilibrium between cerebral blood flow and metabolism is unlikely ^40^. Despite additional cellular effects of ketamine ^41^, we could not discern any impact on electrocorticography when comparing pre-, post-EPCC and awake human brain recordings ^8^.

Since there was no significant change in blood composition or any other appreciable brain parameters under EPCC, any metabolic changes would be subtle enough to not interfere with the ECoG or depth electrode recordings. These recordings are sensitive to metabolism, as attested by extensive experimental studies that document the sensitivity of human and mouse EEG or mouse depth recordings to metabolism and their rapid and steep modulation by energy metabolism state ^29, 31, 42^. Importantly, we used a ECoG and depth electrode recording method that allows for the precise recording of gamma activity, which is exquisitely sensitive to metabolic dysfunction since mitochondrial dysfunction first impairs inhibitory neurons and these neurons are associated with gamma ECoG activity ^29–31, 42^.

## Abbreviations

CBF: cerebral blood flow
CDO_2_: cerebral delivery of oxygen
CVR: cerebral vascular resistance
dH_2_O: deionized water
EPCC: extracorporeal pulsatile circulatory control
LFB: luxol fast blue
MAP: mean arterial pressure
PAS: periodic acid Schiff
RPM: revolutions per minute

## Acknowledgements

We thank the Clark-Samson Laboratory, UT Southwestern Medical Center Department of Neurological Surgery for providing instruments. We also thank Angela Guillory and Cadie Larson for technical assistance with pig maintenance, preparation, and monitoring and Gauri Kathote, Brian Bruscato and Jesus Luna for advice with neurophysiological and circulatory recordings and analyses. The generous contribution of brain probes by Raumedic is also acknowledged. We also thank Ad-Tech Medical for providing sample electrodes and connectors. The illustrations of pig anatomy were modified from biorender.com. The perfusion circuit schematic was created using diagrams.net. The writing of this article was generously supported by an unrestricted gift from the Madeleine Rebekah Verges Foundation to UT Southwestern Medical Center (JMP).

## Author contributions

Conceptualization: JMP, BM, MEJ, MP

Data acquisition: All authors

Methodology: JMP, BM, MP, MEJ, MS, AD, OA

Writing – original draft: AD, MS, BM, MP, JMP

Writing – review & editing: All authors

## Data availability

All data presented or analyzed are available from the corresponding author.

