## Supplement for "Maintenance of pig brain function under extracorporeal pulsatile circulatory control (EPCC)"

**extracorporeal pulsatile circulatory control**

1 The Erik Jonsson School of Engineering and Computer Science, The University of Texas at Dallas, Richardson, Texas, 75080, USA; 2 Rare Brain Disorders Program, Department of Neurology, 3 Department of Cardiovascular & Thoracic Surgery, 4 Department of Neurological Surgery, 5 Animal Resource Center, 6 Department of Internal Medicine, 7 Department of Pathology, 8 Department of Anesthesiology & Pain Management, 9 Department of Physiology, 10 Department of Pediatrics, 11 Eugene McDermott Center for Human Growth & Development/ Center for Human Genetics, The University of Texas Southwestern Medical Center, Dallas, Texas 75390, USA.

¶ Equal contribution

* Present address: Department of Neurosurgery, Loma Linda University Medical Center, Loma Linda, CA, 92354, USA

** Present address: Heart and Vascular Center Brigham and Women's Hospital, Boston, MA 02115, USA

**Date revised**:	May 28, 2023

*** **Correspondence**:

Juan M. Pascual, MD, PhD

The Once Upon a Time Foundation Professor in Pediatric Neurologic Diseases

Ed and Sue Rose Distinguished Professor in Neurology

Director, Rare Brain Disorders Program

UT Southwestern Medical Center

5323 Harry Hines Blvd. Mail code 8813, Dallas, TX 75390-8813

**SUPPLEMENTARY TABLES**

**Supplementary table 1**.

**Blood chemistry under EPCC, standard analysis**. Blood analyses performed in a clinical veterinary facility in subject 2 under native (immediately pre-EPCC) and during EPCC conditions, which included 4 samples obtained hourly.

| Analyte | Native | EPCC | |  |
| --- | --- | --- | --- | --- |
|  |  | Mean | SD | Units |
| Total protein | 5.1 | 4.7 | 0.6 | g/dL |
| Albumin (A) | 2.1 | 2 | 0.3 | g/dL |
| Globulin (G) | 3 | 2.7 | 0.3 | g/dL |
| A/G ratio | 0.7 | 0.75 | 0.06 |  |
| AST (SGOT) | 31 | 36.5 | 3.7 | IU/L |
| ALT (SGPT) | 50 | 46.8 | 6.3 | IU/L |
| Alkaline phosphatase | 90 | 76.8 | 9.0 | IU/L |
| Creatinine | 1.3 | 1.5 | 0.1 | mg/dL |
| Phosphorus | 9 | 10.8 | 2.1 | mg/dL |
| Glucose | 67 | 65.5 | 8.4 | mg/dL |
| Calcium | 9 | 8.3 | 0.5 | mg/dL |
| Sodium | 142 | 146.5 | 3.9 | mEq/L |
| Potassium | 3.9 | 4.7 | 0.7 | mEq/L |
| Chloride | 97 | 90 | 3.4 | mEq/L |
| Cholesterol | 69 | 59.5 | 8.7 | mEq/L |

**Supplementary table 2**.

**Cellular blood composition**. Complete blood counts were obtained in a clinical veterinary facility in subjects 1 and 2 during EPCC and shown in comparison with native value from subject 2. 4 samples were obtained hourly for subject 1 and 3 samples obtained every 1.5 hours for subject 2 and averaged. Standard hematological abbreviations are used.

| Count or index | Native | EPCC | |  |
| --- | --- | --- | --- | --- |
|  |  | Mean | SD | Units |
| WBC | 13.3 | 9.3 | 1.9 | 10^3^/µL |
| RBC | 5.5 | 5.5 | 0.8 | 10^3^/µL |
| HGB | 8.6 | 8.7 | 1.2 | g/dL |
| HCT | 26 | 27.4 | 4.3 | % |
| MCV | 48 | 50.1 | 4.4 | fL |
| MCH | 15.8 | 16.1 | 0.6 | pg |
| MCHC | 33 | 32.4 | 1.9 | g/dL |
| Platelets | 101 | 83.5 | 69.6 | 10^3^/µL |
| Neutrophils | 8512 | 4077 | 844 | /µL |
| Lymphocytes | 3990 | 4885 | 1530 | /µL |
| Monocytes | 266 | 151 | 119 | /µL |
| Eosinophils | 532 | 222 | 138 | /µL |

**Supplementary table 3**.

**Blood chemistry under EPCC, bedside analysis**. Blood analyses were performed with an I-stat analyzer in subject 2 under native (immediately pre-EPCC) and during EPCC conditions, which included 3 samples obtained every 1.5 hours.

| Parameter | Native | Perfusate | |
| --- | --- | --- | --- |
|  |  | Mean | SD |
| pH | 7.64 | 7.47 | 0.07 |
| PCO^2^ (mmHg) | 26.2 | 36.1 | 2.1 |
| PO_2_ (mmHg) | 451 | 365 | 91 |
| HCO_3_ (mmol/L) | 28.1 | 26.4 | 2.8 |
| TCO_2_ (mmol/L) | 29 | 27.7 | 2.5 |
| Saturation O_2_ (%) | 100 | 100 | 0 |
| Na (mmol/L) | 136 | 129 | 3.5 |
| K (mmol/L) | 3.4 | 4.6 | 0.2 |
| Ionized Ca (mmol/L) | 0.54 | 0.34 | 0.01 |
| Glucose (mg/dL) | 291 | 340 | 55 |
| Hematocrit (%) | 19 | 20.3 | 6.5 |
| Hemoglobin (g/dL) | 6.5 | 5.3 | 4.8 |
